# Protein resonance assignment by solid-state NMR based on ^1^H-detected ^13^C-based double-quantum spectroscopy at fast MAS

**DOI:** 10.1101/2021.05.19.444855

**Authors:** Alons Lends, Mélanie Berbon, Birgit Habenstein, Yusuke Nishiyama, Antoine Loquet

## Abstract

Solid-state NMR spectroscopy is a powerful technique to study insoluble and non-crystalline proteins and protein complexes at atomic resolution. The development of proton (^1^H) detection at fast magic-angle spinning (MAS) has considerably increased the analytical capabilities of the technique, enabling the acquisition of ^1^H-detected fingerprint experiments in few hours. Here an approach based on double-quantum (DQ) ^13^C spectroscopy, detected on ^1^H, is introduced at fast MAS (70 kHz) to perform the sequential assignment of insoluble proteins of small size, without any specific deuteration requirement. By combining two three-dimensional ^1^H detected experiments correlating a ^13^C DQ dimension respectively to its intra-residue and sequential ^15^N-^1^H pairs, a sequential walk through DQ (Cα+CO) resonance is obtained. Our approach takes advantage of fast MAS to achieve an efficient sensitivity and the addition of a DQ dimension provides spectral features useful for the resonance assignment process.

## Introduction

Solid-state nuclear magnetic resonance spectroscopy (SSNMR) is a powerful technique to study insoluble, aggregated or non-crystalline biomaterials, ranging from biopolymers ((Kelly et al. 2020), (Zhao et al. 2020),(Goldberga et al. 2018)), carbohydrates ((El Hariri El Nokab and van der Wel 2020)), RNA (Ahmed et al. 2020) or membranes (Dufourc 2021),(Mallikarjunaiah et al. 2019) to larger systems such as protein complexes(Demers et al. 2018) large proteins (Vasa et al. 2018)(Schütz 2021)), protein-ligand interaction ((Vasa et al. 2019)(Elkins and Hong 2019),(Medeiros-Silva et al. 2019)), misfolded proteins (König et al. 2019), amyloid (Tycko 2016)fibrils ((Jaroniec 2019),(Loquet et al. 2018)), helical filaments (Habenstein et al. 2019), viruses (Lecoq et al. 2020), (Gupta et al. 2020), (Lu et al. 2020)), membrane proteins (McDermott 2009)(Tang et al. 2013)(Mandala et al. 2018) or whole cells ((Narasimhan et al. 2020)). In the two past decades, structural investigation of biomolecules at atomic resolution by SSNMR has made dramatic analytical improvements with the introduction of direct proton (^1^H) detection(Ishii et al. 2001)(Reif et al. 2001)(Paulson et al. 2003)(Zhou et al. 2009) combined with the use of magic-angle spinning (MAS) probes operating at fast frequencies(Nishiyama 2016)(Böckmann et al. 2015)(Cala-De Paepe et al. 2017)(Sternberg et al. 2018)(Xue et al. 2018)(Ishii et al. 2018)(Schledorn et al. 2020), nowadays commercially available at MAS frequency of 60-110 kHz. ^1^H-detection takes advantage of the high gyromagnetic ratio of proton spins leading to an excellent sensitivity(Ishii et al. 2001)(Demers et al. 2011), together with the fact that ^1^H are highly abundant in biomolecules. Fast MAS probes require sub-milligram sample quantities, and well-resolved 3D and 4D ^1^H-detected experiments of deuterated ((Knight et al. 2011)(Agarwal et al. 2014)(Fricke et al. 2017)) and fully protonated proteins ((Stanek et al. 2016)(Andreas et al. 2016)(Xiang et al. 2015)(Vasa et al. 2019) have been obtained for protein samples.

Two-dimensional (2D) ^1^H-detected correlation experiments such as 2D ^13^C-^1^H spectra offer a powerful spectroscopic tool to monitor chemical shift perturbations. These spectral “fingerprints” have the potential to become the new routine experiment for SSNMR investigations of proteins, in analogy to the ^15^N-^1^H HSQC experiment for solution NMR, as we recently demonstrated to rapidly compare amyloid fibril conformation of various α-synuclein fibril polymorphs based on 2D ^15^N/^13^C-^1^H spectra (De Giorgi et al. 2020). Although few minutes to hours are required to obtain these spectral fingerprints, a tremendous time is still required to perform the so-called resonance assignment step, usually based on 3D experiments to establish triple resonance connectivity’s between ^1^H, ^13^C and ^15^N atoms(Barbet-Massin et al. 2014)(Penzel et al. 2015)(Fricke et al. 2017). The assignment process relies on two main steps: first the identification of intra-residue signals including backbone and side chain atoms, and then the establishment of one or more sequential connectivity’s between two adjacent residues in the sequence. These sequential assignment experiments have been designed to sequentially link intra (*i*) and proceeding (*i-1*) Cα, CO and Cβ nuclei in correlation with their ^15^N and ^1^H^N^ backbone nuclei. The efficiency of these experiments(Barbet-Massin et al. 2014) typically relies on the MAS frequency, the protonation level and the structural homogeneity of the samples. For protonated samples at 100 kHz MAS, the backbone assignment can be extended with the set of 6 Hα-detected experiments(Stanek et al. 2016). The combination of both amide H^N^ and CαHα-detected experiments can significantly simplify the sequential assignment procedure(Wiegand et al. 2020)(Schubeis et al. 2020), even more by using H^N^ / CαHα simultaneous acquisition (Sharma et al. 2020)(Stanek et al. 2020) as recently developed for automatic resonance assignment.

Several spectroscopic and biochemistry solutions have been developed to overcome the resonance assignment process, such as the use of higher spectral dimensionality (>3), however, it requires more experimental time in order to achieve a sufficient signal-to-noise ratio (SNR)(Xiang et al. 2014)(Fraga et al. 2017)(Zinke et al. 2017). Another solution is the use of non-uniform isotope labeling strategies to reduce the number of ^1^H and ^13^C sites, such as sparse(Higman et al. 2009), selective(Hoffmann et al. 2018) and segmental labeling schemes(Skrisovska et al. 2010). These methods can reduce the number of overlapping peaks, however additional steps of sample preparation are required, associated with extra costs.

The uniqueness of ^13^C resonances (e.g. Cα of Gly, Cβ for Ala, Thr and Ser) offers one of the most valuable starting point for the identification of the residue-type, and another spectroscopic option to help the resonance assignment process is to decrease the peak overlap via the acquisition of the ^13^C dimension in the double-quantum (DQ) mode. To overcome peak overlapping, a better spectral dispersion can be created from the summation of different spectral regions, ideally in protein samples with Cα+Cβ and Cα+CO. Addition of a DQ dimension helps to increase the spectral dispersion of otherwise overlapping peaks(Luca and Baldus 2002)(Koźmiński and Zhukov 2004) in protein NMR spectra. The DQ dimension is achieved when the magnetization is excited and then back recovered to a zero quantum (SQ) mode using recoupling schemes. This approach has been implemented for dipolar- and scalar-based 2D(Lesage et al. 1997)(Hong 1999) and 3D spectra INADEQUATE-type experiments have been used to simplify the analysis of ^13^C-^13^C correlations(Luca and Baldus 2002)(Chordia et al. 2021). Li *et al*(Li et al. 2020) recently reported the detection of DQ-SQ correlations with a ^13^C detection to increase the spectral resolution of side-chain carboxyl groups of Glu, Asp, Gln and Asn residues. Xiao et al (Xiao et al. 2021) applied Cα-Cβ DQ filtering for Ala, Thr and Ser residues to simplify the 2D and 3D ^15^N-^13^C-^13^CX experiments. We (Tolchard et al. 2018) and others(Ward et al. 2011) took advantage of direct ^1^H detection to generate a ^13^C DQ dimension leading to an efficient polarization transfer for correlating ^13^C to ^1^H spins in protein samples. In 2011, Ladizhansky and coworkers reported the use of 3D DQ(CXCa)NH and DQ(CaCO)NH experiments at moderate MAS frequency (28 kHz) using a SPC-5 sequence on a deuterated membrane protein sample to help the backbone assignment process(Ward et al. 2011). In 2018 we demonstrated, in the case of a fully protonated protein amyloid fibril sample, that a 3D DQ(CXCY)CYH correlation experiment at 70 kHz MAS can provide an efficient strategy to detect side-chains ^1^H and ^13^C resonances (Tolchard et al. 2018). Because a single 3D experiment was developed in this previous study, the work only permitted a signal identification without the possibility to perform sequential assignment.

In the following, we demonstrate the complete sequential resonance assignment of a medium-size protein assembly through the use of two DQ experiments to access intra-residue and sequential correlations, without a specific deuteration requirement. We take advantage of several spectroscopic features obtained at fast MAS (70 kHz) combined with the use of ^1^H-detection to implement high sensitivity 3D correlation experiments using a ^13^C DQ dimension. Combination of DQ(Ca_i_CX_i_)N_i_H_i_ and DQ(CO_i-1_Ca_i-1_)N_i_H_i_ experiments, complemented by a simple 3D CαNH, leads to a straightforward assignment process based on sequential (Cα+CO) plans. Our approach is demonstrated with the assignment of the amyloid fibrils HET-s in its fully protonated form.

## Materials and methods

### Sample preparation

^13^C/^15^N-labeled, fully protonated HET-s(218-289) was expressed in E. coli, purified and aggregated based on previously described protocols (Balguerie et al. 2003). Approximately 800 μg of HET-s(218-289) amyloid fibrils were introduced in a 1 mm JEOL SSNMR rotor. The tripeptide N-formyl-L-Met-L-Leu-L-Phe (fMLF) was purchased from CortecNet and packed into a 1 mm JEOL ssNMR rotor.

### Solid-state NMR spectroscopy

All spectra were recorded with a 21.1 T (900 MHz ^1^H Larmor frequency) JEOL spectrometer equipped with a 1 mm triple resonance HCN MAS probe. The sample temperature was kept at 290 K and the MAS frequency set to 70 kHz. For the BaBa excitation and recovery, we used 1.33 μs long pulses at 188 kHz for total of 457 μs. The detailed spectral acquisition and processing parameters are provided in **Tables S2** and **S3**. The spectra were processed using JEOL Delta and NMRPipe (Delaglio et al. 1995) software and analyzed using CcpNmr program (Vranken et al. 2005).

## Results and discussion

### Pulse sequence design

Inspired by the work of Ladizhansky (Ward et al. 2011) and our work (Tolchard et al. 2018), we propose two 3D SSNMR MAS experiments (the overall sequence shown in **Fig.1A**.) using DQ excitation and recovery using the back-to-back (BaBa)-xy16 recoupling(Sommer et al. 1995) (Feike et al. 1996), considering that such recoupling is efficient at fast (>60 kHz) MAS regime(Saalwächter et al. 2011)(Tolchard et al. 2018). Moreover, the broadband performance of the BaBa coupling is proportional to the MAS frequency, making it particularly attractive under high B_0_ field at faster spinning frequencies. It is also robust to radio-frequency field inhomogeneity(Saalwächter et al. 2011) and provides high DQ filtering efficiency. During the BaBa recoupling the magnetization is transferred only between spatially proximate ^13^C spins via dipolar couplings due to dipolar truncation, since recoupled DQ Hamiltonians do not commute each other. Thus, the transfer will preferentially occur between covalently coupled ^13^C spins, making the BaBa recoupling a highly selective way to achieve polarization transfer between two carbon sites within the same residue. We implemented BaBa recoupling in (i) a DQ(Ca_i_CX_i_)N_i_H_i_ sequence, which is detecting the ^13^C DQ dimension as the sum of (Cα+Cβ) and (Cα+CO) of intra residue and (ii) a DQ(CO_i-1_Ca_i-1_)N_i_H_i_, experiment which displays the sum of (Cα+CO) of the proceeding residue.

**Fig. 1.**
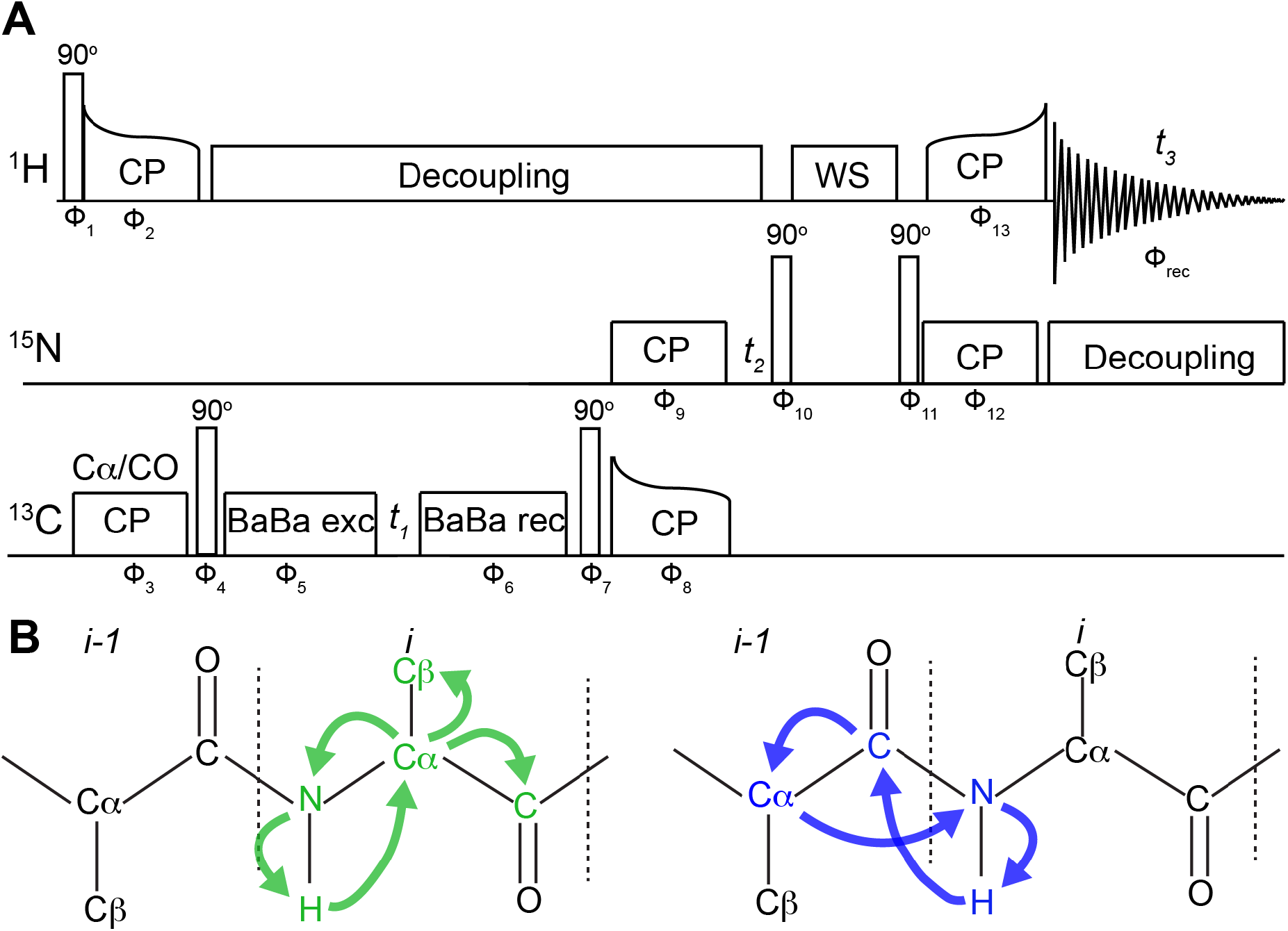
**A** Pulse sequence of the 3D DQ(Ca_i_CX_i_)N_i_H_i_, DQ(CO_i-1_Ca_i-1_)N_i_H_i_ experiments. The initiation of corresponding polarization transfers are achieved through the carrier value of the first ^1^H-^13^C cross-polarization transfer. The applied phase cycling was executed with the following scheme Φ_1_={8(x), 8(-x)}, Φ_2_={x},Φ_3_={x}, Φ_4_={-x}, Φ_5_={x,-x,y,-y}, Φ_6_={x}, Φ_7_={-x}, Φ_8_={x}, Φ_9_={4(x), 4(y)},Φ_10_={x}Φ_11_={-y}, Φ_12_={16(x), 16(y)}, Φ_13_={x}, Φ_rec_={2(x,y),2(y,x),2(y,x),2(x,y),2(y,x),2(x,y),2(x,y),2(y,x)}. **B** Schematic representation of the polarization transfer pathways and involved spins for DQ(Ca_i_CX_i_)N_i_H_i_ (green) and DQ(CO_i-1_Ca_i-1_)N_i_H_i_ (blue) experiments.

The overall structure for the DQ(Ca_i_CX_i_)N_i_H_i_ and DQ(CO_i-1_Ca_i-1_)N_i_H_i_ 3D SSNMR pulse sequences with schematic magnetization pathways are shown in **Fig.1B**. Both sequences start with a cross-polarization (CP) transfer from ^1^H spins to either Cα or CO spins, depending on the placement of the frequency carrier, to provide a frequency-selective polarization on Cα and CO for the DQ(Ca_i_CX_i_)N_i_H_i_ and DQ(CO_i-1_Ca_i-1_)N_i_H_i_ experiments respectively. The following 90° pulse is applied to create the longitudinal magnetization. The DQ coherence for the first dimension detection (t1) is generated using a BaBa-XY16 excitation(Saalwächter et al. 2011). During this period, the signal can be acquired as the sum of (C+C) resonances, following two polarization patterns. In the DQ(Ca_i_CX_i_)N_i_H_i_ experiment, a DQ(Cα_i_CX_i_) is generated to access the sum of Cα to both carbonyl (CO) and Cβ nuclei, in order to probe the intra-residue spin system. In the DQ(CO_i-1_Ca_i-1_)N_i_H_i_experiment, a DQ(COCα) is generated to provide the sum of CO to the adjacent Cα. The reconversion is achieved through another block of BaBa recoupling. The 10 kHz CW ^1^H decoupling was applied during each BaBa 90° pulse and t1 and t2 acquisition periods. The subsequent ^13^C 90° pulse sets the magnetization back to transverse plane and it is transferred to ^15^N spins through a ^13^C-^15^N CP step, followed by the second dimension detection (t2). The next ^15^N 90° pulse sets the magnetization along longitudinal plane for the MISSISIPPI(Zhou and Rienstra 2008) water suppression and another 90^o 15^N pulse transfers the magnetization back to the transverse plane. In the last step, a ^15^N-^1^H CP step is applied to transfer the magnetization back to ^1^H spins for the direct detection (t3), applying a WALTZ16 decoupling for ^15^N spins. The overall transfer steps can be written as 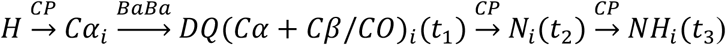 for 3D DQ(Ca_i_CX_i_)N_i_H_i_ and as 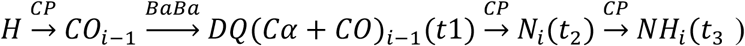 for 3D DQ(CO_i-1_Ca_i-1_)N_i_H_i_ experiments. To obtain a selective polarization transfer to the adjacent carbon, a short mixing was chosen (229 µs) during the BaBa excitation. In this way, a selective DQ coherence is created to correlate Cα_i_ to its neighboring Cβ_i_ and CO_i_ spins in the DQ(Ca_i_CX_i_)N_i_H_i_ experiment. In the DQ(CO_i-1_Ca_i-1_)N_i_H_i_ experiment, the carbonyl CO_i-1_ has only one neighboring carbon (i.e. Cα_i-_1). Both sequences were implemented in the same way, with only difference being the position of the frequency carrier for the first ^1^H-^13^C and second ^13^C-^15^N CP transfer. Hence the optimized condition can be conveniently transferred between two sequences.

### Polarization transfer efficiency at fast MAS

In order to estimate the polarization transfer efficiency of BaBa recoupling under 70 kHz MAS, we compared 1D ^1^H spectra after a (^1^H-Cα)-^1^H and a (^1^H-Cα-BaBa-Cα)-^1^H polarization pathway respectively. We used the fully protonated protein HET-s(218-289) as a protein benchmark. HET-s(218-289) forms amyloid fibrils, this sample was already used by our laboratory to test pulse sequences at fast MAS regime(Stanek et al. 2016)(Tolchard et al. 2018). From the overlay of spectra showed in **Fig.S1**, we determine that the DQ filter yields to 40% of intensity compared to the pulse sequence employing only CP transfers. This result is comparable to values obtained for other SSNMR homonuclear recoupling sequences (De Pape 2012).

Next, we acquired 1D ^1^H spectra to evaluate the transfer efficiencies of DQ(Ca_i_CX_i_)N_i_H_i_ and DQ(Ca_i_CX_i_)N_i_H_i_ experiments for the fully protonated fMLF sample and HET-s(218-289) amyloid fibrils. We compared the two DQ-based sequence to well-established hCaNH, hCONH hcoCacoNH, hCOcaNH and hcaCbcaNH experiments (described in details in (Barbet-Massin et al. 2014). To set up a reference for intensity comparison, we used a 1D hNH experiment. This experiment uses only two CP steps and was used before as the standard for the comparison of ^1^H detected spectra(Barbet-Massin et al. 2014)(Andreas et al. 2015)(Xiang et al. 2015)(Vallet et al. 2020). The relative intensities of all experiments to the hNH experiment are summarized in **Table S1**. DQ(Ca_i_CX_i_)N_i_H_i_ and DQ(Ca_i_CX_i_)N_i_H_i_ experiments compared to hCaNH, hCOcaNH, hcaCbcaNH and hCONH experiment are summarized in **Table 1**. Each spectrum was adjusted based on the number of scans. As already observed for deuterated samples(Barbet-Massin et al. 2014)(Xiang et al. 2015)(Penzel et al. 2015) the highest intensities is observed for the hCaNH experiment, leading to ∼0.17 transfer efficiency compared to the hNH experiment for the fMLF sample. For deuterated back-exchanged protein samples at 40-60 kHz MAS, these values were higher, i.e. ∼0.20-0.30 for hCaNH (Barbet-Massin et al. 2014)(Fricke et al. 2017)(Vallet et al. 2020). These differences can be attributed to longer ^13^C and ^1^H *T*_*1ρ*_ relaxation times for deuterated samples, leading to more efficient transfers and a higher proton sensitivity. Here, the use of the DQ filter leads to a remarkable polarization transfer efficiency compared to the well-established SQ experiments (**Table 1**), demonstrating the benefit of adding the BaBa excitation and reconversion recoupling schemes. The overall DQ(Ca_i_CX_i_)N_i_H_i_ and DQ(CO_i-1_Ca_i-1_)N_i_H_i_ experiments lead to a transfer efficiency of ∼1.4-5% compared to the hNH experiment for both samples (**Table S1**). The DQ(Ca_i_CX_i_)N_i_H_i_ transfer efficiency to correlate intra-residue spins is ∼0.31 compared to the hCaNH experiment for the fMLF sample. A similar result (∼0.37) is obtained for HET-s(218-289) amyloid fibrils (**Table 1**). For the DQ experiment correlating CO_i-1_ spins to its sequential N_i_H_i_ pair, a polarization transfer efficiency compared to the hCONH experiment of 0.16 and 0.19 was measured for fMLF and HET-s(218-289) samples respectively (**Table 1**). For inter-residual Ca correlation DQ(CO_i-1_Ca_i-1_)N_i_H_i_ experiment is 0.25 and 0.44 less intensive compare to the hcoCacoNH sequence. The efficiency of the DQ(Ca_i_CX_i_)N_i_H_i_ to detect intra-residue CO is higher to the hCOcaNH sequence (**Table S1**), which is important experiment for sequential assignment, but is one of the least efficient of 3D assignment experiments (Barbet-Massin et al. 2014)(Penzel et al. 2015). Thus, obtaining intra-residue CO chemical shift with a higher sensitivity is additional advantage for DQ(Ca_i_CX_i_)N_i_H_i_ sequence over INEPT based experiment. To estimate the efficiency of the DQ(Ca_i_CX_i_)N_i_H_i_ sequence to provide a polarization transfer to intra-residue Cβ spins, we compared to the hcaCbcaNH sequence, which has been reported as one of the most sensitive sequence to detect Cβ spins(Penzel et al. 2015). The relative intensity of the hcaCbcaNH experiment was ∼1% compared to a hNH experiment for HET-s(218-289) fibrils (∼3% for fMLF). A relative intensity of ∼5% for fMLF and HET-s(218-289) was obtained for the DQ(Cα_i_CX_i_)N_i_H_i_ sequence (**Table S1**). These values obtained here for protonated systems are lower to reported values obtained for deuterated proteins (6%-16%)(Barbet-Massin et al. 2014)(Penzel et al. 2015)(Vallet et al. 2020), due to the shorter *T*_*2*_*’* transverse relaxation times for protonated sample. The lower absolute sensitivity for hCOcaNH and hcaCbcaNH experiments is attributed to the relatively poor sensitivity of scalar-based CC transfers with multiple echo delays. However, this drawback is compensated by a more selective polarization transfer, as hCOcaNH and hcaCbcaNH are optimized to selectively target CO and Cβ spins respectively, while in DQ-based experiments a broader polarization transfer is obtained.

**Table 1.** Relative intensity of 1D experiments recorded on fMLF and HET-s(218-289) amyloid fibrils at 70 kHz MAS.

| Experiment comparison | Relative intensity |  |
| --- | --- | --- |
|  | fMLF | HET-s(218-289) |
| DQ(C$\alpha_i$CX$_i$)N$_i$H$_i$ / hCaNH | 0.31 | 0.37 |
| DQ(CO$_{i-1}$Ca$_{i-1}$)N$_i$H$_i$ / hCONH | 0.16 | 0.19 |
| DQ(C$\alpha_i$CX$_i$)N$_i$H$_i$ / hCOcaNH | 4.2 | 3 |
| DQ(CO$_{i-1}$Ca$_{i-1}$)N$_i$H$_i$ / hcoCacoNH | 0.25 | 0.44 |
| DQ(C$\alpha_i$CX$_i$)N$_i$H$_i$ / hcaCbcaNH | 1.7 | 4.8 |

### Protein resonance assignment strategy using ^13^C DQ-detected experiments

The resonance assignment strategy relies on the establishment of connectivities between intra-residue pairs (Cα-CO and Cα-Cβ) to their NH pair and between the (Cα-CO) pair to its sequential NH pair. To disentangle the SQ Cα chemical shift to the signal provided by the (Cα+Cβ) and (Cα+CO) DQ resonances, a single 3D hCaNH experiment is additionally required. In the 3D DQ(Ca_i_CX_i_)N_i_H_i_ spectrum, each individual NH strip displays the sum of intra-residue (Cα+Cβ) and (Cα+CO) resonances. The Cβ(*i*) and CO(*i*) shifts can be calculated by subtracting Cα (*i*) shifts (acquired in hCaNH spectrum) from the additions of (Cα+Cβ) or (Cα+CO) DQ resonances. In this manner we can obtain the Cα, Cβ and CO triplet for the same residue. Then the 3D DQ(CO_i-1_Ca_i-1_)N_i_H_i_ experiment provides the sum of (Cα+CO) resonance of the proceeding *i-1* residue and is matched to the sum of (Cα+CO) of intra-residue resonance. As a consequence, the difference of (Cα_i_+CO_i_) and (Cα_i_+CO_i_) resonances will result in a single Cα_i-1_ shift. In such manner, a sequential backbone walk is achieved by employing only three 3D experiments, which then give access to intra-residue H, N, Cα, Cβ and CO as well as Cα and CO chemical shifts of the proceeding residue.

### Completeness of the assignment strategy

For the fMLF tripeptide, we assign all anticipated intra-residue and sequential correlations in the two 3D DQ-detected spectra (the 2D ^1^H-^13^C strips from 3D spectra are shown in **Fig.2A**). The assignment was confirmed with comparison of previously published chemical shifts for the fMLF peptide(Struppe et al. 2017). For the Cβ assignments in 2D strips of hcaCbcaNH spectrum we found 2 out for 3 expected peaks. The absence of Cβ peak for the Leu residue (**Fig.2A**), might be attributed to ongoing dynamics at different time scale, which can hinder the scalar coupling-based magnetization transfer. Leu Cβ spin is detected in the 3D DQ(Ca_i_CX_i_)N_i_H_i_ spectrum. Due to even lower sensitivity of 1D hcaCbcaNH experiment for HET-s(218-289) we didn’t acquired as 3D spectrum. For HET-s(218-289) the Cβ shifts were obtained solely from the DQ(Ca_i_CX_i_)N_i_H_i_ spectrum by subtraction of Cα shifts from the sum of (Cα+Cβ) peaks.

**Fig. 2.**
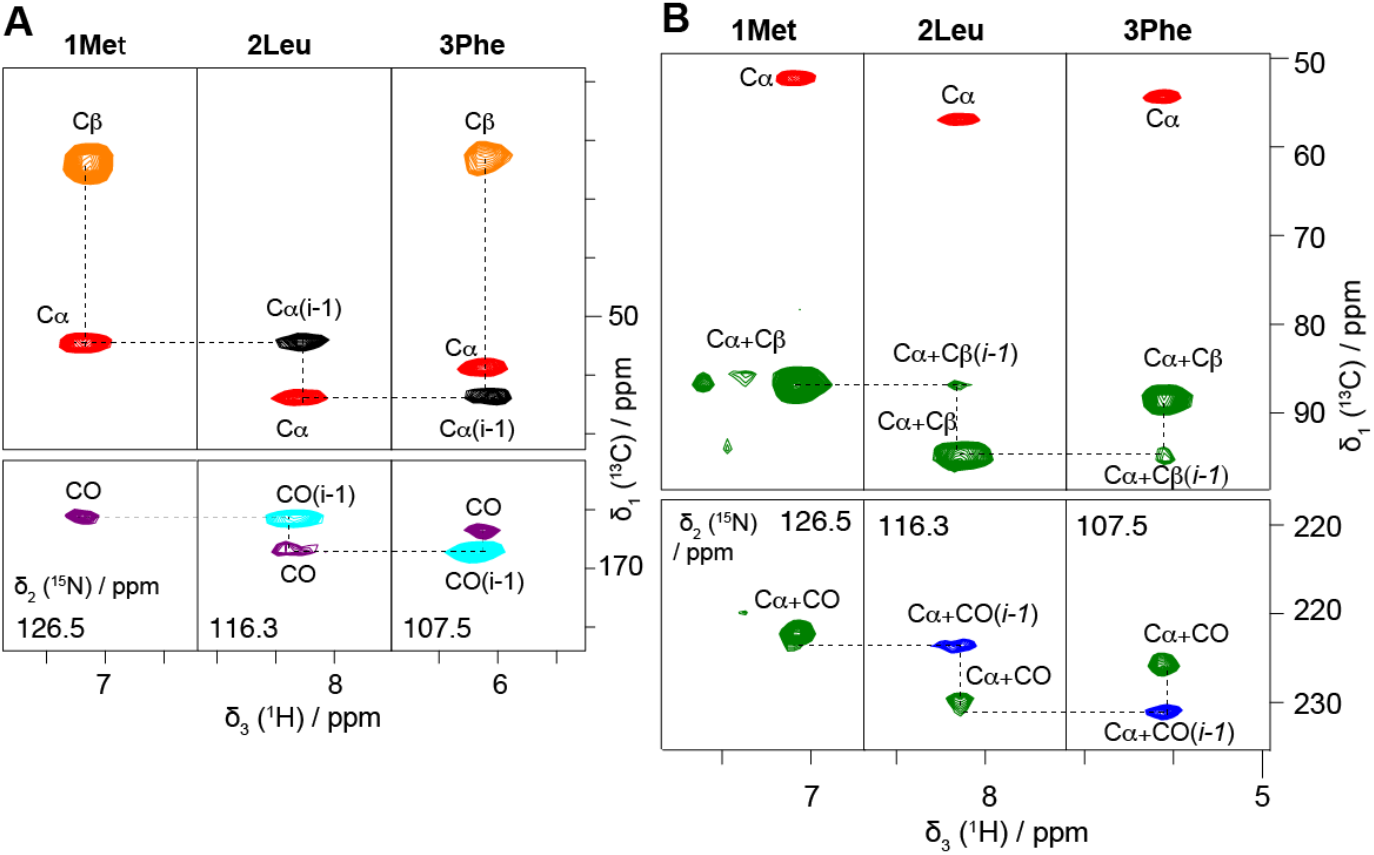
2D projections of ^1^H-detected SSNMR experiments recorded on fMLF. **A** the 2D ^1^H-^13^C strips along ^15^N dimension for the 3D hCaNH (red), hCONH (cyan), hcoCacoNH (black), hcaCbcaNH (orange) and hCOcaNH (purple) spectra. **B** The 2D ^1^H-^13^C strips along ^15^N dimension for the hCaNH (red), DQ(Ca_i_CX_i_)N_i_H_i_ (green) and DQ(CO_i-1_Ca_i-1_)N_i_H_i_ (blue) spectra, the sequential backbone walk is shown with dotted lines.

Both DQ(Ca_i_CX_i_)N_i_H_i_ and DQ(CO_i-1_Ca_i-1_)N_i_H_i_ experiments also provide a larger ^13^C connection maps compare to SQ ^13^C dimension experiments. The sequential backbone walk using ^13^C DQ experiments were conducted through 2D ^1^H-^13^C projections along ^15^N dimension of 3D DQ(Ca_i_CX_i_)N_i_H_i_ and DQ(CO_i-1_Cα_i-1_)N_i_H_i_ spectra in a combination with 3D hCaNH spectrum (**Fig.2B**). We use the 3D DQ(Ca_i_CX_i_)N_i_H_i_ spectrum to access (Cα+Cβ) and (Cα+CO) intra-residue correlations. In the 3D DQ(CO_i-1_Cα_i-1_)N_i_H_i_ experiment, the sum (Cα+CO) of *i-1* peaks (blue peaks in **Fig.2B**) can be directly connected to (Cα+CO) of *i* residue (green peaks in **Fig.2B**). Interestingly, we noticed additional weak intensity peaks in 2D planes of 3D DQ(Ca_i_CX_i_)N_i_H_i_. These peaks perfectly matched the sum (Cα+Cβ) of the proceedin*g* residue. Similar observation based on the observation of weak *i-1* correlations has been described for solution NMR HNCa experiment(Kay et al. 1990). The origins of these unexpected peaks might be due to non-selective band excitation of BaBa recoupling (**Fig.2B**). As a consequence, the DQ coherence can spread to weakly coupled carbons through dipolar couplings. On one side, these additional peaks are reducing the peak intensities of intra-residue correlations due to dipolar truncation. On the other side, they provide additional information to establish a sequential link. Overall the DQ(Ca_i_CX_i_)N_i_H_i_ and DQ(CO_i-1_Ca_i-1_)N_i_H_i_ spectra showed expected intra-residue and sequential correlation for the fMLF peptide.

Next, we demonstrated the approach on a medium-size insoluble protein assembly, the amyloid fibrils of HET-s(218-289). At 70 kHz MAS, we measured a ^1^H line-width of ∼250 Hz on a 21 Tesla magnet (900 MHz ^1^H Larmor frequency) from a 2D hNH spectrum (**Fig.S2**). Such a ^1^H line-width is close to the value reported for the same fibrillar protein system acquired at 100 kHz MAS (at 1 GHz ^1^H Larmor frequency) (Stanek et al. 2016). We observed slightly lower signal intensities (compare to hNH) for 3D INEPT-based assignment experiments for HET-s(218-289) amyloid fibrils compared to the fMLF sample, due to additional relaxation effects that lead to less efficient polarization transfers. However, the overall intensity for the DQ(Ca_i_CX_i_)N_i_H_i_ experiment was comparable to fMLF and it was even higher for the DQ(CO_i-1_Ca_i-1_)N_i_H_i_ experiment. It might indicate that both ^13^C DQ-based experiments are more prone to slower transverse relaxation times as present for the HET-s sample. An overlay of 2D DQ-^13^C/^15^N projections for 3D DQ(Ca_i_CX_i_)N_i_H_i_ and hCaNH experiments (**Fig.3A**) and DQ(CO_i-1_Ca_i-1_)N_i_H_i_ and hCONH experiments (**Fig.3B**). All cross-peaks in the 3D DQ-detected experiments are well spread out, and the inclusion of the Cβ resonances does not increase the spectral overlap since it is encoded via a DQ dimension. As a consequence, most (Cα+Cβ) resonances are spread around 85-130 ppm, and do not overlap with Cα signals (45-65 ppm). The lowest sum of (Cα+Cβ) peaks arise from Ala residues, hence they are shifted in the upfield part of the spectrum, around 70 ppm (**Fig.3A**). Gly residues, due to the absence of a Cβ atom, appear only as the sum of (Cα+CO) peaks (**Fig.3B**) shifted upfield (∼ 45 ppm) compare to other DQ resonances. As expected, the most downfield shifted signals in (Cα+Cβ) region belong to Thr and Ser residues, due to their higher chemical shift values. All other residues have DQ resonances that appear in between the sum of (Cα+Cβ) resonances of Thr/Ser and Ala residues. We notice that for several aliphatic residues, resonances overlap in the hCaNH spectrum (red spectrum in **Fig.3A**), for instance residues 267V, 231I and 229K. This can cause additional ambiguities for the protein chemical shift assignment. In the DQ(Ca_i_CX_i_)N_i_H_i_ spectrum (green spectrum in **Fig.3A**) the DQ-detected resonance for the same residues were more clearly dispersed due to generated sum of (Cα+Cβ) resonance in the DQ ^13^C dimension. This spectral feature of DQ ^13^C-detected experiments provides a similar advantage for the detection of CO_i-1_ resonances in DQ(CO_i-1_Ca_i-1_)N_i_H_i_ compared to hCONH experiment (spectral overlay is shown in **Fig.3B**). We employed a similar strategy, as previously described for the fMLF peptide, to carry out the sequential assignment of HET-s(218-289) resonances. The resonance assignment strategy on a protein sample is illustrated in **Fig.3C**. The 2D ^1^H-DQ^13^C strip plots along ^15^N dimension are particularly efficient to provide a sequential protein backbone assignment using combination of 3D DQ(Ca_i_CX_i_)N_i_H_i_, DQ(CO_i-1_Ca_i-1_)N_i_H_i_ and hCaNH experiments. Using this approach, we achieve the assignment of 87.5 % of backbone HN, 81.2 % of intra-residue ^13^C and we establish a sequential connectivity for 87.5% of the residues in the rigid core of HET-s(218-289) amyloid fibrils.

**Fig. 3.**
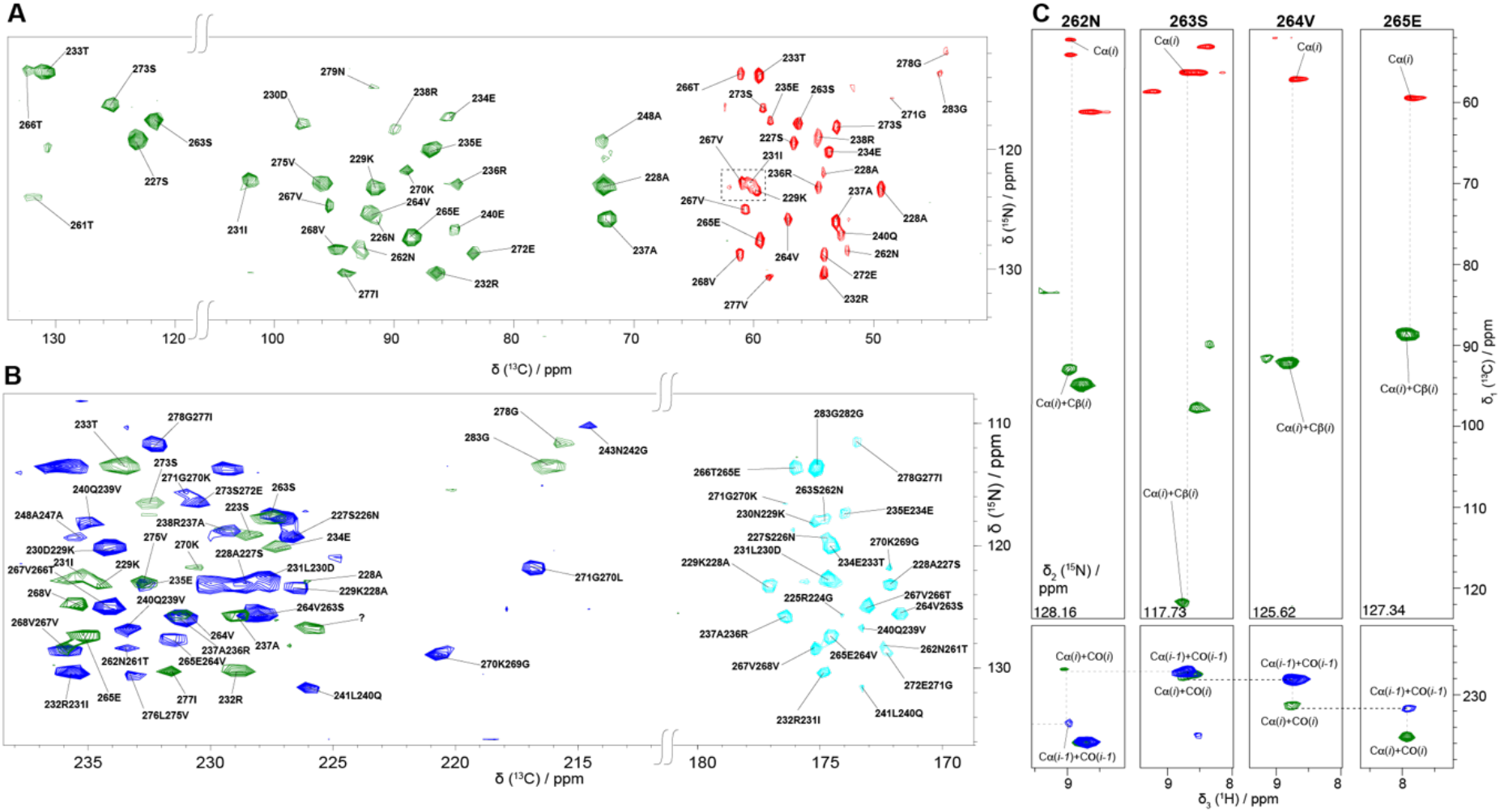
2D projections of ^1^H-detected SSNMR experiments recorded on HET-s(218-289) amyloid fibrils. **A** 2D ^13^C-^15^N projections of 3D DQ(Ca_i_CX_i_)N_i_H_i_ (green) and hCaNH (red) experiments, showing intra-residue Cα, CO, Cα+Cβ and Cα+CO correlations. The overlapping peaks in the N-Cα region is indicated with the dotted square. **B** 2D ^13^C-^15^N projections of DQ(CO_i-1_Ca_i-1_)N_i_H_i_ (blue) and hCONH (cyan) experiments, showing the inter residue CO and Cα+CO regions. **C** 2D ^1^H-^13^C strips along ^15^N dimension for the sequential backbone walk is shown with dotted lines.

One advantage of the DQ-based sequential backbone assignment is the requirement of a set of only three 3D experiments. The two 3D experiments (DQ(Ca_i_CX_i_)N_i_H_i_ and DQ(CO_i-1_Ca_i-1_)N_i_H_i_) were recorded in 5 days and provided enough sensitivity and spectral dispersion to perform a *de novo* assignment (in combination with a hCaNH, which took 15 h to acquired) of fully protonated HET-s(218-289) amyloid fibrils. The combination of both 3D DQ(Ca_i_CX_i_)N_i_H_i_ and DQ(CO_i-1_Ca_i-1_)N_i_H_i_ spectra with hCaNH allows to identify backbone chemical shifts, to link sequential Cα-CO pairs and to detect intra-residue Cβ spins. Additionally, the approach artificially increases the ^13^C signal dispersion for otherwise overlapping peaks due to the DQ dimension. In order to cover the large chemical shift differences in the DQ ^13^C dimension, both DQ(Ca_i_CX_i_)N_i_H_i_ and DQ(CO_i-1_Ca_i-1_)N_i_H_i_ experiments need however to be acquired with a sufficiently large sampling rates in the indirect DQ ^13^C dimension (details shown in **Tables S2** and **S3**). In that regard, the number of scans for an adequate measurement time must be leveraged in order to achieve high enough SNR.

Completeness of the HET-s(218-289) resonance assignment using the DQ-detected approach was determined to be ∼87 % of ^13^C, ^15^N and ^1^H resonances, considering the rigid residues of the amyloid core observable in a CP experiment. This result is comparable to previously reported resonance assignment strategies on the same protein using conventional ^13^C-detected (Siemer et al. 2006b)(Van Melckebeke et al. 2010) and ^1^H-detected SSNMR experiments (Smith et al. 2017) (Tolchard et al. 2018) (Stanek et al. 2016). The missing residues in ^1^H detected spectra have been reported for other proteins(Torosyan et al. 2019)(Jirasko et al. 2021). We hypothesize that these atoms are involved in a faster dynamic range that averages out dipole couplings and therefore weakens the CP transfer step. The access to these residues would be possible using INEPT or direct polarization during the first magnetization initiation step as demonstrated by Meier and coworkers(Lange et al. 2011)(Siemer et al. 2006a)(Siemer 2020).

## Conclusions

In this study, we demonstrate the potential of introducing a ^13^C DQ dimension in 3D SSNMR ^1^H-detected experiments at fast MAS to perform protein resonance assignment. The use of a ^13^C DQ dimension using the Baba recoupling provides a remarkable sensitivity to achieve ^13^C-^13^C polarization transfer at fast MAS (70 kHz) and increases the ^13^C spectral dispersion. The approach was demonstrated on fully protonated HET-s fibrils to access the detection of ^13^C, ^15^N and ^1^H backbone spins, detect intra-residue Cβ, Cα, CO and sequentially link Cα-CO pairs. For small-to-medium size proteins typically studied by SSNMR (i.e. ∼30-100 amino-acids detected in a CP experiment), the approach is a promising alternative to conventional out-and-back techniques using CC scalar transfers popularized by Pintacuda and coworkers(Barbet-Massin et al. 2013)(Barbet-Massin et al. 2014). The ever-increasing faster MAS frequencies in combination with higher magnetic fields will be also beneficial for DQ-based sequences, due to increased broadband range for the BaBa recoupling and longer ^13^C and ^1^H *T’*_*2*_ relaxation times. Further on, we are going to expand ^13^C DQ sequences to combine them with Hα detection methods and implement them into automatic assignment programs.

## Acknowledgments

We acknowledge financial support from the European Research Council (ERC) under the European Unions Horizon 2020 research and innovation program (ERC-2015-StG GA no. 639020 to A. Loquet). The project was supported by FranceAgriMer and the CNIV through the program PNDV (project ATOMIVINE n° 297772) and the IdEx Bordeaux (Chaire d’Installation to B.H., ANR-10-IDEX-03-02). A. Lends was supported by the Swiss National Science Foundation for early postdoc mobility project P2EZP2_184258. This work was supported by JSPS KAKENHI Grant Number 20K05483 and in part by the JST-Mirai Program (Grant No. JPMJMI17A2, Japan) to Y. Nishiyama.

