## Supplementary Information for "Protein resonance assignment by solid-state NMR based on ^1^H-detected ^13^C-based double-quantum spectroscopy at fast MAS"

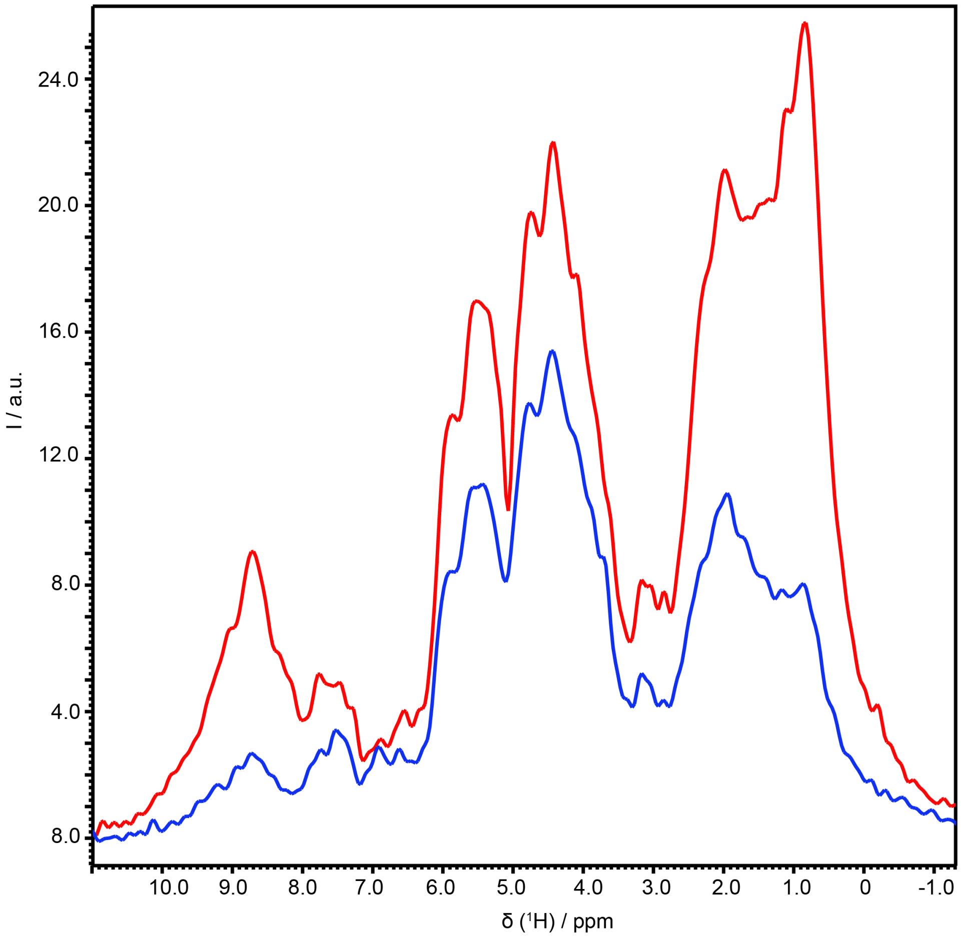


**Fig.S1**. The overlay of 1D spectra for HET-s(218-289) amyloid fibrils, fully protonated, recorded at 21 T, 70 kHz MAS frequency, for 16 scans. In red: ($^{1}$H-Cα)-$^{1}$H spectrum. In blue: ($^{1}$H-Cα-BaBa-Cα)-$^{1}$H.

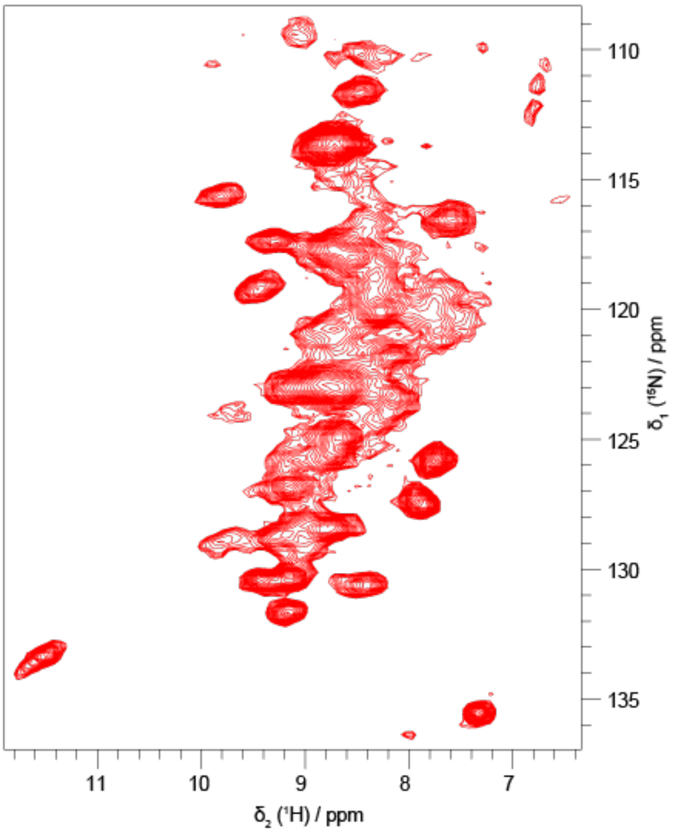


**Fig.S2**. 2D hNH spectrum of fully protonated HET-s(218-289) amyloid fibrils, recorded at 21 T, 70 kHz MAS frequency.

**Table S1**. The relative intensity from 1D compare to hNH spectrum at 70 kHz MAS.

| Experiment | Relative intensity, % | |
|---|---|---|
| | **fMLF** | **HET-s(218-289)** |
| hCaNH | 17 | 12.7 |
| hCONH | 8.5 | 16.0 |
| hcoCacoNH | 5.5 | 6.7 |
| hCOcaNH | 1.2 | 1.5 |
| hcaCbcaNH | 3.0 | 1.0 |
| DQ($Ca_iCX_i$)$N_iH_i$ | 5 | 4.8 |
| DQ($CO_{i-1}Ca_{i-1}$)$N_iH_i$ | 1.4 | 3.0 |

**Table S2**. The SSNMR acquisition parameters for fMLF. All spectra were acquired at 21 Tesla magnet, 70 kHz MAS frequency. The States-TTPI acquisition mode was applied for indirect dimensions. For a comparison of the 1D spectra the same acquisition parameters were used.

| Experiment | 3D hCaNH | 3D hCONH | 3D hcoCacoNH | 3D hCOcaNH | 3D hcaCbcaNH | 3D h($Ca_iCx_i$)$N_iH_i$ | 3D h($CO_{i-1}Ca_{i-1}$)$N_iH_i$ |
|---|---|---|---|---|---|---|---|
| Transfer 1 | HC-CP | HC-CP | HC-CP | HC-CP | HC-CP | HC-CP | HC-CP |
| $^1$H field/kHz | 48 | 48 | 48 | 48 | 48 | 48 | 48 |
| $^{13}$C field/kHz | 2-38 | 2-38 | 2-38 | 2-38 | 2-38 | 2-38 | 2-38 |
| Shape | ramp $^{13}$C | ramp $^{13}$C | ramp $^{13}$C | ramp $^{13}$C | ramp $^{13}$C | ramp $^{13}$C | ramp $^{13}$C |
| Time/ms | 2 | 2 | 2 | 2 | 1 | 2 | 1.5 |
| $^{13}$C carrier/ppm | 55 | 171.5 | 55 | 171.5 | 40 | 80 | 171.5 |
| Transfer 2 | CaN-CP | CON-CP | CC INEPT | CC INEPT | CC INEPT | BaBa | BaBa |
| $^{15}$N field/kHz | 55.6-36.0 | 55.6-36.0 | | | | | |
| $^{13}$C field/kHz | 9.6 | 9.6 | | | | | |
| Shape | ramp $^{15}$N | ramp $^{15}$N | | | | | |
| Time/ms | 16 | 16 | 5 | 5 | 5 | 0.457 | 0.457 |
| $^1$H decoupling | WALTZ16 | WALTZ16 | | | | | |
| $^1$H decoupling field/ kHz | 10 | 10 | | | | | |
| Transfer 3 | NH-CP | NH-CP | | | | | |
| $^1$H field/kHz | 18.6 | 18.6 | | | | | |
| $^{15}$N field/kHz | 36.0-55.6 | 36.0-55.6 | | | | | |
| Shape | ramp $^{15}$N | ramp $^{15}$N | | | | | |
| Time/ms | 0.5 | 0.5 | | | | | |
| Transfer 4 | | | NH-CP | NH-CP | NH-CP | NH-CP | NH-CP |
| $^1$H field/kHz | | | 18.6 | 18.6 | 18.6 | 18.6 | 18.6 |
| $^{15}$N field/kHz | | | 36.0-55.6 | 36.0-55.6 | 36.0-55.6 | 36.0-55.6 | 36.0-55.6 |
| Shape | | | ramp $^{15}$N | ramp $^{15}$N | ramp $^{15}$N | ramp $^{15}$N | ramp $^{15}$N |
| Time/ms | | | 0.5 | 0.5 | 0.5 | 0.5 | 0.5 |
| t1 increments | 8 | 8 | 8 | 8 | 8 | 128 | 16 |
| Windows function | QSine3 | QSine3 | QSine3 | QSine3 | QSine3 | QSine3 | QSine3 |
| Sweep width (t1)/kHz | 2.3 | 2.3 | 2.3 | 2.3 | 4.5 | 70 | 3.5 |
| Acquisition time (t1)/ms | 3.5 | 3.5 | 3.5 | 3.5 | 1.8 | 1.8 | 4.5 |
| $^1$H decoupling | WALTZ16 | WALTZ16 | WALTZ16 | WALTZ16 | WALTZ16 | WALTZ16 | WALTZ16 |
| $^1$H decoupling field/ kHz | 10 | 10 | 10 | 10 | 10 | 10 | 10 |
| t2 increments | 8 | 8 | 8 | 8 | 8 | 8 | 8 |
| Windows function | QSine 3 | QSine 3 | QSine 3 | QSine 3 | QSine 3 | QSine 3 | QSine 3 |
| Sweep width (t2)/kHz | 2.7 | 2.7 | 2.7 | 2.7 | 2.7 | 2.7 | 2.7 |
| Acquisition time (t2)/ms | 2.9 | 2.9 | 2.9 | 2.9 | 2.9 | 2.9 | 2.9 |
| $^{15}$N/$^{13}$C decoupling | WALTZ16 | WALTZ16 | WALTZ16 | WALTZ16 | WALTZ16 | WALTZ16 | WALTZ16 |
| $^{15}$N/$^{13}$C decoupling field/ kHz | 10 | 10 | 10 | 10 | 10 | 10 | 10 |
| t3 increments | 1024 | 1024 | 1024 | 1024 | 1024 | 1024 | 1024 |
| Windows function | QSine 3 | QSine 3 | QSine 3 | QSine 3 | QSine 3 | QSine 3 | QSine 3 |
| Sweep width (t1)/kHz | 125 | 125 | 125 | 125 | 125 | 125 | 125 |
| Acquisition time (t1)/ms | 8.2 | 8.2 | 8.2 | 8.2 | 8.2 | 8.2 | 8.2 |
| Interscan delay/s | 2 | 2 | 2 | 2 | 2 | 2 | 2 |
| Number of scans | 2 | 2 | 2 | 18 | 16 | 16 | 16 |
| Measurement time | 18 min | 18 min | 18 min | 2.9 h | 2.6 h | 40 h | 5 h |

**Table S3**. The SSNMR acquisition parameters for HET-s(218-289). All spectra were acquired at 21 Tesla magnet, 70 kHz MAS frequency. The States-TTPI acquisition mode was applied for indirect dimensions. For a comparison of the 1D spectra the same acquisition parameters were used.

| Experiment | 2D hNH | 3D hCaNH | 3D hCONH | 3D h($Ca_iCx_i$)$N_iH_i$ | 3D h($CO_{i-1}Ca_{i-1}$)$N_iH_i$ |
|---|---|---|---|---|---|
| **Transfer 1** | HN-CP | HC-CP | HC-CP | HC-CP | HC-CP |
| **$^1$H field/kHz** | 57 | 58.2-84 | 58.2-84 | 20 | 20 |
| **$^{13}$C field/kHz** | | 11 | 11 | 82-42 | 82-42 |
| **$^{15}$N field/kHz** | 13.6-28.9 | | | | |
| **Shape** | ramp $^{15}$N | ramp $^1$H | ramp $^1$H | ramp $^{13}$C | ramp $^{13}$C |
| **Time/ms** | 1 | 0.5 | 4 | 0.5 | 2 |
| **$^{13}$C carrier/ppm** | | 55 | 171.5 | 100 | 112.5 |
| **Transfer 2** | NH-CP | CaN-CP | CON-CP | BaBa | BaBa |
| **$^1$H field/kHz** | 57 | | | | |
| **$^{15}$N field/kHz** | 28.9-13.6 | 70.9-51.2 | 70.9-51.2 | | |
| **$^{13}$C field/kHz** | | 9 | 9 | | |
| **Shape** | ramp $^{15}$N | ramp $^{15}$N | ramp $^{15}$N | | |
| **Time/ms** | 0.4 | 16 | 16 | 0.457 | 0.457 |
| **$^1$H decoupling** | WALTZ16 | WALTZ16 | WALTZ16 | WALTZ16 | WALTZ16 |
| **$^1$H decoupling field/ kHz** | 10 | 10 | 10 | 10 | 10 |
| **Transfer 3** | | NH-CP | NH-CP | CaN-CP | CON-CP |
| **$^1$H field/kHz** | | 18.6 | 18.6 | | |
| **$^{13}$C field/kHz** | | | | 23.6- 11.6 | 9.6 |
| **$^{15}$N field/kHz** | | 44.7-64.3 | 44.7-64.3 | 53 | 68.7-49 |
| **Shape** | | ramp $^{15}$N | ramp $^{15}$N | ramp $^{13}$C | ramp $^{15}$N |
| **Time/ms** | | 0.5 | 0.5 | 10 | 16 |
| **Transfer 4** | | | | NH-CP | NH-CP |
| **$^1$H field/kHz** | | | | 68 | 18.6 |
| **$^{15}$N field/kHz** | | | | 28.9-13.6 | 44.7-64.3 |
| **Shape** | | | | ramp $^{15}$N | ramp $^{15}$N |
| **Time/ms** | | | | 0.4 | 0.5 |
| **t1 increments** | 128 | 44 | 16 | 32 | 24 |
| **Windows function** | QSine 3 | QSine 3 | QSine 3 | QSine 3 | QSine 3 |
| **Sweep width (t1)/kHz** | 4.56 | 6.7 | 2 | 70 | 7 |
| **Acquisition time (t1)/ms** | 28.1 | 6.6 | 15.7 | 0.46 | 3.4 |
| **$^1$H decoupling** | WALTZ16 | WALTZ16 | WALTZ16 | WALTZ16 | WALTZ16 |
| **$^1$H decoupling field/ kHz** | 10 | 10 | 10 | 10 | 10 |
| **t2 increments** | 1024 | 32 | 32 | 32 | 32 |
| **Windows function** | QSine 3 | QSine 3 | QSine 3 | QSine 3 | QSine 3 |
| **Sweep width (t2)/kHz** | 125 | 3.1 | 3.1 | 3.1 | 3.1 |
| **Acquisition time (t2)/ms** | 8.2 | 10.3 | 10.3 | 10.3 | 10.3 |
| **$^{15}$N/$^{13}$C decoupling** | WALTZ16 | WALTZ16 | WALTZ16 | WALTZ16 | WALTZ16 |
| **$^{15}$N/$^{13}$C decoupling field/ kHz** | 10 | 10 | 10 | 10 | 10 |
| **t3 increments** | | 1024 | 1024 | 1024 | 1024 |
| **Windows function** | | QSine 3 | QSine 3 | QSine 3 | QSine 3 |
| **Sweep width (t3)/kHz** | | 125 | 125 | 125 | 125 |
| **Acquisition time (t3)/ms** | | 8.2 | 8.2 | 8.2 | 8.2 |
| **Interscan delay/s** | 1.5 | 1 | 2 | 1.3 | 0.95 |
| **Number of scans** | 32 | 8 | 8 | 32 | 64 |
| **Measurement time/h** | 4 | 15.5 | 5.6 | 55.5 | 64.5 |
